# Deciphering the genetic diversity of landraces with high-throughput SNP genotyping of DNA bulks: methodology and application to the maize 50k array

**DOI:** 10.1101/2020.05.19.103655

**Authors:** Mariangela Arca, Tristan Mary-Huard, Brigitte Gouesnard, Aurélie Bérard, Cyril Bauland, Valérie Combes, Delphine Madur, Alain Charcosset, Stéphane D. Nicolas

## Abstract

Genebanks harbor original landraces carrying many original favorable alleles for mitigating biotic and abiotic stresses. Their genetic diversity remains however poorly characterized due to their large within genetic diversity. We developed a high-throughput, cheap and labor saving DNA bulk approach based on SNP Illumina Infinium HD array to genotype landraces. Samples were gathered for each landrace by mixing equal weights from young leaves, from which DNA was extracted. We then estimated allelic frequencies in each DNA bulk based on fluorescent intensity ratio (FIR) between two alleles at each SNP using a two step-approach. We first tested either whether the DNA bulk was monomorphic or polymorphic according to the two FIR distributions of individuals homozygous for allele A or B, respectively. If the DNA bulk was polymorphic, we estimated its allelic frequency by using a predictive equation calibrated on FIR from DNA bulks with known allelic frequencies. Our approach: (i) gives accurate allelic frequency estimations that are highly reproducible across laboratories, (ii) protects against false detection of allele fixation within landraces. We estimated allelic frequencies of 23,412 SNPs in 156 landraces representing American and European maize diversity. Modified Roger’s genetic Distance between 156 landraces estimated from 23,412 SNPs and 17 SSRs using the same DNA bulks were highly correlated, suggesting that the ascertainment bias is low. Our approach is affordable, easy to implement and does not require specific bioinformatics support and laboratory equipment, and therefore should be highly relevant for large-scale characterization of genebanks for a wide range of species.

## INTRODUCTION

Genetic resources maintained *in situ* or *ex situ* in genebanks represent a vast reservoir of traits/alleles for future genetic progress and an insurance against unforeseen threats to agricultural production (Tanksley 1997; Hoisington *et al*. 1999; Kilian and Graner 2012; McCouch *et al*. 2012). Due to their local adaptation to various agro-climatic conditions and human uses, landraces are particularly relevant to address climate change and the requirements of low input agriculture (Fernie *et al*. 2006; McCouch *et al*. 2012; Mascher *et al*. 2019). For instance, maize displays considerable genetic variability, but less than 5 % of this variability has been exploited in elite breeding pools, according to (Hoisington *et al*. 1999). However, landraces are used to a very limited extent, if any, in modern plant breeding programs, because they are poorly characterized, genetically heterogeneous and exhibit poor agronomic performance compared to elite material (Kilian and Graner 2012; Strigens *et al*. 2013; Brauner *et al*. 2019; Mascher *et al*. 2019; Hölker *et al*. 2019). Use of molecular techniques for characterizing genetic diversity of landraces and their relation with the elite germplasm is essential for a better management and preservation of genetic resources and for a more efficient use of these germplasms in breeding programs (Hoisington *et al*. 1999; Mascher *et al*. 2019).

The genetic diversity of landraces conserved *ex situ* or *in situ* has been extensively studied using various types of molecular markers such as restriction fragment length polymorphism (RFLP) or simple sequence repeat (SSR) in maize (Dubreuil and Charcosset 1998; Dubreuil *et al*. 1999; Rebourg *et al*. 1999, 2001; Gauthier *et al*. 2002; Rebourg *et al*. 2003; Reif *et al*. 2005b; Vigouroux *et al*. 2005; Reif *et al*. 2005a; Camus-Kulandaivelu 2006; Dubreuil *et al*. 2006; Eschholz *et al*. 2010; Mir *et al*. 2013), in Pearl Millet (Bhattacharjee *et al*. 2002), cabbage (Dias *et al*. 1991; Divaret *et al*. 1999), Barley (Parzies *et al*. 2000; Backes *et al*. 2003; Hagenblad *et al*. 2012), pea (Hagenblad *et al*. 2012), oat (Hagenblad *et al*. 2012), rice (Ford-Lloyd *et al*. 2001; McCouch *et al*. 2012), Alfalfa (Pupilli *et al*. 2000; Segovia-Lerma *et al*. 2003) and fonio millet (Adoukonou-Sagbadja *et al*. 2007). SSRs have proven to be markers of choice for analyzing diversity in different plant species and breeding research, because of their variability, ease of use, accessibility of detection and reproducibility (Liu *et al*. 2003; Reif *et al*. 2006; Yang *et al*. 2011). Nevertheless, the development of SSR markers requires a substantial investment of time and money. Allele coding is also difficult to standardize across genotyping platforms and laboratories, a major drawback in a genetic resources characterization context. SNPs have become the marker of choice for various crop species such as maize (Ganal *et al*. 2011), rice (McCouch *et al*. 2010), barley (Moragues *et al*. 2010) and soybean (Lam *et al*. 2010). They are the most abundant class of sequence variation in the genome, co-dominantly inherited, genetically stable and appropriate to high-throughput automated analysis (Rafalski 2002). For instance, maize arrays with approx. 50,000 and 600,000 SNP markers are available since 2010 (Illumina MaizeSNP50 array, Ganal *et al*. 2011) and 2013 (600K Affymetrix Axiom, Unterseer et al., 2013), respectively. SNP arrays may however lead to some ascertainment bias notably when diversity analysis was performed on a plant germplasm distantly related from those that have been used to discover SNP put on the array (Nielsen 2004; Clark *et al*. 2005; Hamblin *et al*. 2007; Inghelandt *et al*. 2011; Frascaroli *et al*. 2013). Properties of SNP array regarding diversity analysis have to be carefully investigated to evaluate ascertainment biais. For maize 50K Infinium SNP array, only “PZE” prefixed SNPs (so called later PZE SNPs in this study) give consistent results for diversity analysis as compared with previous studies based on SSR markers and are therefore suitable for assessing genetic variability (Inghelandt *et al*. 2011; Ganal *et al*. 2011; Bouchet *et al*. 2013; Frascaroli *et al*. 2013). 50K Infinium SNP array has been used successfully to decipher genetic diversity of inbred lines (van Heerwaarden *et al*. 2011; Bouchet *et al*. 2013; Frascaroli *et al*. 2013; Rincent *et al*. 2014), landraces using either doubled haploids (Strigens *et al*. 2013) or a single individual per accession (van Heerwaarden *et al*. 2011; Arteaga *et al*. 2016), or teosinte with few individuals per accession (Aguirre-Liguori *et al*. 2017).

Due to high diversity within accessions, characterization of landraces from allogamous species such as maize or alfalfa should be performed based on representative sets of individuals (Reyes-Valdés *et al*. 2013, Segovia-Lerma et al., 2002, Dubreuil and Charcosset., 1998). Despite the recent technical advances, genotyping large numbers of individuals remains very expensive for many research groups. To bring costs down, estimating allele frequencies from pooled genomic DNA (also called “allelotyping”) has been suggested as a convenient alternative to individual genotyping using high-throughput SNP arrays (Sham *et al*. 2002; Teumer *et al*. 2013) or using Next Generation Sequencing (Schlötterer *et al*. 2014). It was successfully used to decipher global genetic diversity within maize landraces using RFLPs (Dubreuil and Charcosset 1998; Dubreuil *et al*. 1999; Rebourg *et al*. 2001, 2003; Gauthier *et al*. 2002) and SSR markers (Reif *et al*. 2005a; Camus-Kulandaivelu 2006; Dubreuil *et al*. 2006; Yao *et al*. 2007; Mir *et al*. 2013). Various methods for estimating gene frequencies in pooled DNA have been developed for RFLP (Dubreuil and Charcosset 1998), SSR (LeDuc *et al*. 1995; Perlin *et al*. 1995; Daniels *et al*. 1998; Lipkin *et al*. 1998; Breen *et al*. 1999) and SNP marker arrays in human and animal species (Hoogendoorn *et al*. 2000; Craig *et al*. 2005; Brohede 2005; Teumer *et al*. 2013; Gautier *et al*. 2013). These methods have been successfully used to detect QTL (Lipkin *et al*. 1998), to decipher genetic diversity (Segovia-Lerma *et al*. 2003; Dubreuil *et al*. 2006; Pervaiz *et al*. 2010; Johnston *et al*. 2013; Ozerov *et al*. 2013), to perform genome wide association studies (Barcellos *et al*. 1997; Sham *et al*. 2002; Baum *et al*. 2007), to identify selective sweep (Elferink *et al*. 2012) or to identify causal mutation in tilling bank (Abe *et al*. 2012). Genotyping DNA bulks of individuals from landraces with SNP arrays could therefore be interesting to characterize and manage genetic diversity in plant germplasm. SNP arrays could be notably a valuable tool to identify selective sweep between landraces depending on their origin, to manage plant germplasm collection at worldwide level (e.g. identify duplicate), to identify landraces poorly used so far in breeding programs or to identify genomic regions where diversity has been lost during the transition from landraces to inbred lines (Arca et al., *in prep*).

In this study, we developed a new DNA bulk method to estimate allelic frequencies at SNPs based on Fluorescent Intensity data produced by the maize 50K Illumina SNP array (Ganal *et al*. 2011). Contrary to previous methods that have been mostly developed for QTL detection purposes, our approach is dedicated to genome-wide diversity analysis in plant germplasm since it protects against false detection of alleles. Additionally, calibration of equations for predicting allelic frequencies of DNA bulks for each SNP is based on controlled pools with variable allelic frequencies rather than only heterozygous genotypes as in previous methods (Hoogendoorn *et al*. 2000; Brohede 2005; Peiris *et al*. 2011; Teumer *et al*. 2013). As a proof of concept, we applied our new approach to maize by estimating allelic frequencies of 23,412 SNPs in 156 maize landraces representative of European and American diversity present in genebanks (Arca et al., *in prep*). To our knowledge, it is the first time that a DNA bulk approach was used on 50K maize high-throughput SNP array to study genetic diversity within maize landraces germplasm.

## RESULTS

We developed a new method to estimate allelic frequencies of SNPs within pools of individuals using the fluorescent intensity ratio (FIR) between A and B alleles from Illumina MaizeSNP50 array. Briefly, allelic frequencies at SNPs belonging to MaizeSNP50 array were estimated within 156 maize landraces by pooling randomly 15 individuals per population and by calibrating a predictive two-step model (Figure 1). We considered only the subsample of 32,788 prefixed PZE markers (so called PZE SNPs) that have proven suitable for diversity analyses (Ganal *et al*. 2011). Among these SNPs, we selected 23,412 SNPs that passed weighted deviation (wd) quality criteria (wd>50). This removed SNPs for which estimated allelic frequency deviated strongly from expected allelic frequency (Figure. S1 A, B, C, D, E, F and G for the threshold choice and validation).

**Figure 1:**
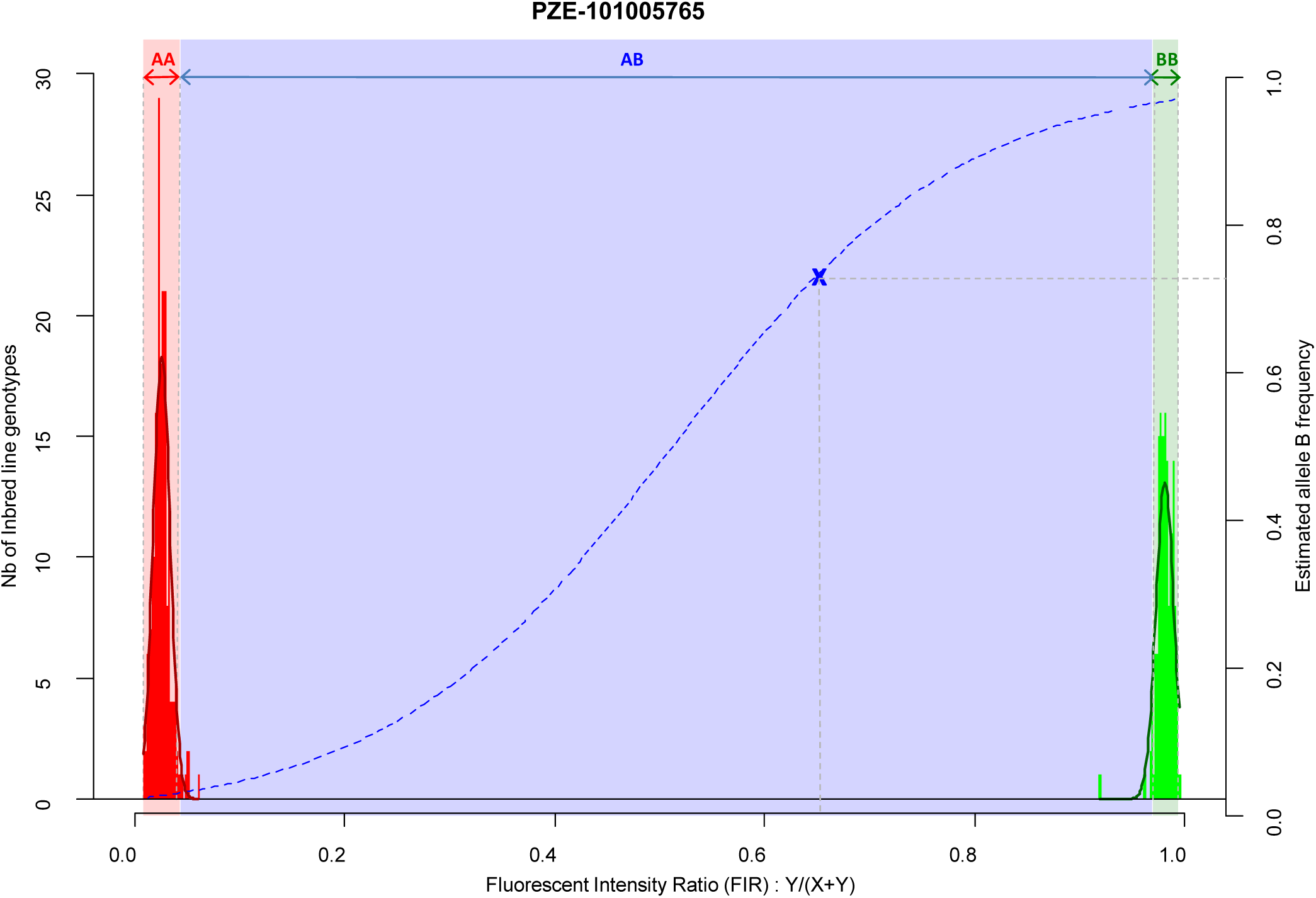
Two-step approach for estimating allelic frequency in DNA pools, exemplified by marker PZE-101005765. Red and green histograms correspond to the fluorescent intensity ratio (FIR) distribution for inbred lines homozygote for allele A (AA) and B (BB), respectively. Red and green curves indicate the corresponding Gaussian distributions. Red, Blue, and Green areas correspond to the FIR for which landraces are declared homozygous for allele A, polymorphic and homozygous for allele B after testing for fixation of alleles A and B. Dotted blue line corresponds to the curve of the logistic regression adjusted on 1,000 SNPs and two series of controlled pools. Blue cross corresponds to a landrace represented by a DNA bulk of 15 individuals, with its observed FIR on X axis and predicted frequency on Y axis.

### Accuracy of allelic frequency prediction and detection of allele fixation

In order to prevent erroneous detection of alleles within landraces, we first tested for each landrace whether allele A or allele B was fixed at a given SNP locus (Figure 1). We tested for each SNP whether the FIR of the landrace was included within one the two Gaussian distributions drawn from mean and variance of FIR of genotypes AA and BB within the inbred line panel (Figure 1). For landraces that were considered polymorphic after this first step (allele fixation rejected for both alleles), we estimated allelic frequency based on FIR by using a unique logistic regression model for the 23,412 SNPs, calibrated with a sample of 1,000 SNPs (Figure 1). Parameters of the logistic model were adjusted on these 1,000 SNPs using FIR of two series of controlled pools whose allelic frequencies were known (Figure S2). We obtained these pools by mixing various proportion of two series of three inbred lines with known genotypes (Table 1). The1,000 SNPs were selected to maximize the allelic frequency range within controlled pools (Table 1). The logistic regression was calibrated on 1,000 SNPs taken together rather than for each SNP individually to avoid the ascertainment bias that would be generated by selecting only SNPs polymorphic in the controlled pools (Figure S3) and to reduce loss of accuracy in prediction for SNPs displaying limited allelic frequency range in two controlled pools (Figure S4). To investigate the loss of accuracy of the prediction curve due to a reduction in allelic frequency ranges in controlled pool, we progressively removed at random from one to 15 samples from the calibration set of the 1000 above described SNPs. The mean absolute error (MAE) between 1000 replications increased significantly from 4.14 % to 8.54 % when removing more samples (Table 2). For comparison, MAE was 7.19 % using a cross-validation approach in which the predictive equation was calibrated with a random subsample of 800 out of 1000 SNPs, and then applied to estimate allelic frequencies for the remaining 200 SNPs (Table S1). Calibrating the logistic regression between FIR and allelic frequency in controlled pool based on 1000 SNPs therefore appears well adapted to prevent ascertainment bias while increasing globally prediction accuracy (Figure S4). Finally, we observed that MAE was higher for balanced allelic frequencies than for almost fixed allelic frequencies (Figure 2 and Tables S2). Accordingly, the dispersion of predicted frequencies were larger for expected allelic frequencies near 0.5 than for fixed alleles (Table S2).

**Table 1:**
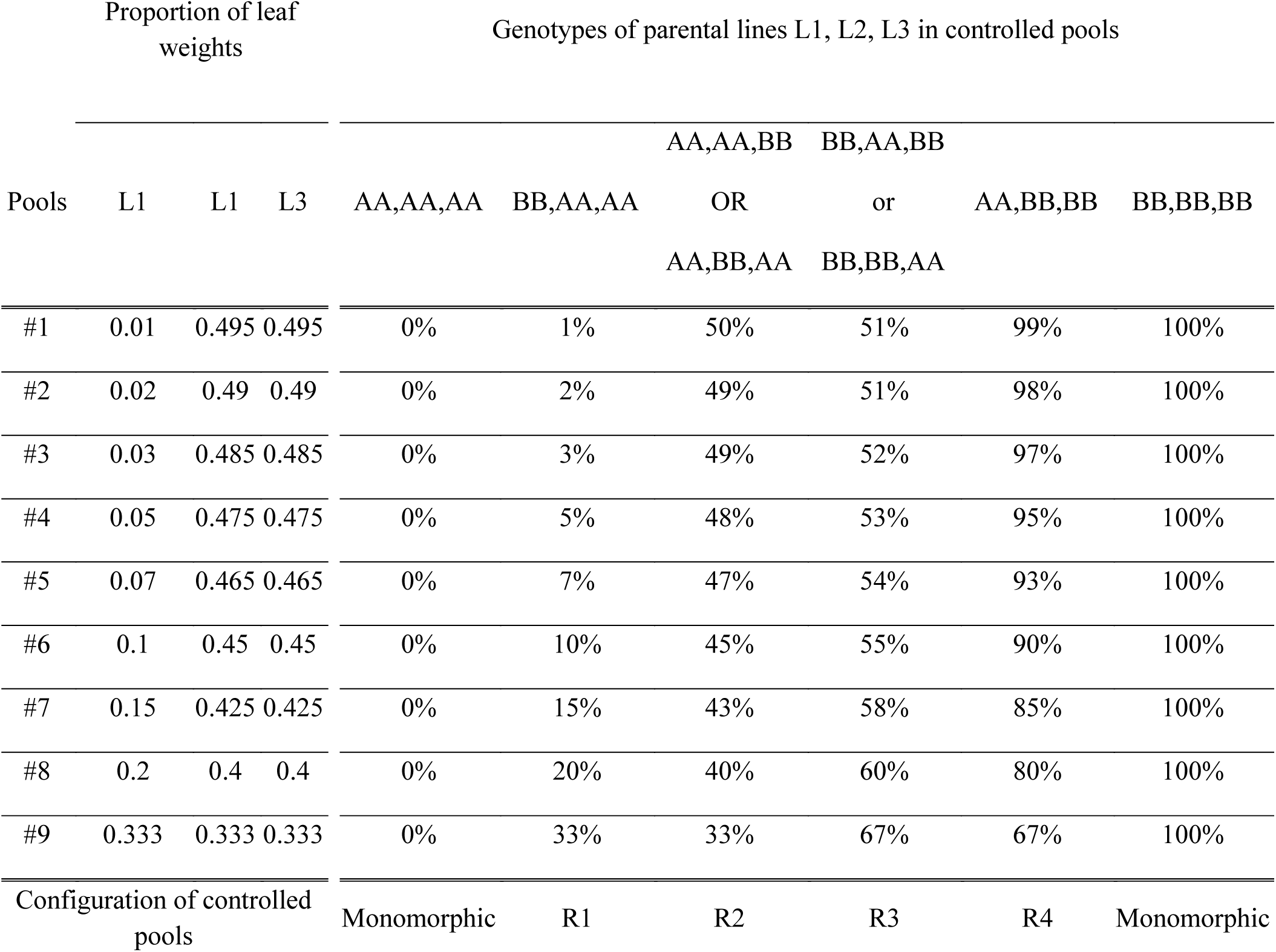
Expected frequencies of allele B for the nine controlled pools obtained by varying the proportions of leaf weights of three inbred lines (L1, L2, L3) according to their genotypes at a bi-allelic SNP coded A/B. Heterozygous genotypes for inbred lines were not considered in this table.

**Table 2:** Mean absolute error (MAE) in frequency estimation for 1,000 SNPs used to calibrate logistic regression equations. MAE is estimated by a cross-validation procedure in which a number of pools comprised between 1 and 15 among 18 is removed at random from the calibration set. This procedure was repeated 1,000 times for each SNP.

| # of removed samples | # of repetitions | Mean absolute error (MAE) |  |
| --- | --- | --- | --- |
|  |  | Mean | SD* |
| 1 | 1000 | 0.0414 | 0.0219 |
| 3 | 1000 | 0.0428 | 0.0226 |
| 5 | 1000 | 0.0447 | 0.0232 |
| 8 | 1000 | 0.0484 | 0.0245 |
| 10 | 1000 | 0.0522 | 0.0257 |
| 12 | 1000 | 0.0582 | 0.0274 |
| 15 | 1000 | 0.0854 | 0.0309 |

\* SD = Standard deviation

**Figure 2:**
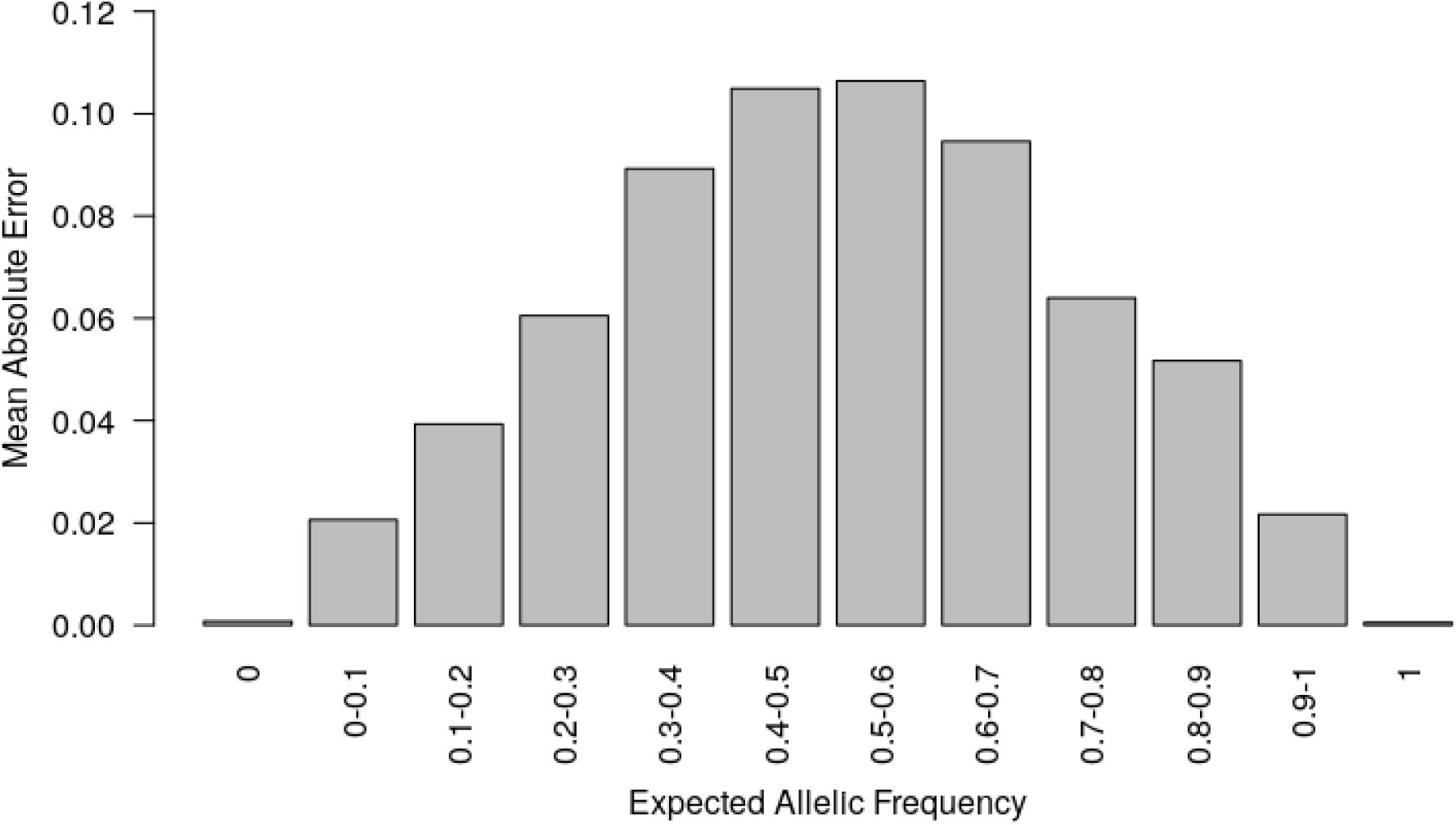
Mean absolute error (MAE) according to the known allelic frequency in two series of controlled pools. MAE measured the absolute difference between allelic frequencies predicted by the two-step approach and those expected from the genotypes of parental lines in two series of controlled pools for 23,412 SNPs. MAE is averaged for each interval of expected allelic frequency across all SNPs.

### Reproducibility of frequency across laboratories and samples

We evaluated the reproducibility of the method across laboratories by comparing FIR of one series of controlled pools from two different laboratories using all PZE SNPs or 23,412 SNPs selected using *wd* criterion (Figure 3). The correlation between the two different laboratories for controlled pools was very high (r^2^>0.98) whether we selected the SNPs based on *wd* criterion or not. Beyond reproducibility across laboratories, the precision of frequency estimation depends on the sampling of individuals within landraces (Table 3). The precision of frequency estimation was addressed both by numerical calculation and by the independent sampling of 15 different individuals (30 different gametes) within 10 landraces (biological replicate). For both numerical calculations and biological replicates, the sampling error was higher for loci with balanced allelic frequencies than for loci that are close to fixation (Table 3, Figure 4). Sampling error also decreased as the number of sampled individuals increased (Table 3). Considering a true frequency of 50% within landraces, we expect that 95% of frequency estimates lie between 31.30% and 68.70% when sampling 15 individuals per landrace and 42.9 to 57.13% when sampling 100 individuals per landrace (Table 3). Considering biological replicates, allelic frequencies of the two biological replicates over 23,412 SNPs were highly correlated except for population Pol3 (Table S3). When excluding Pol3, 94.5% of points were located within the 95% confidence limits accounting for the effect of sampling alone, suggesting that the error inherent to the frequency estimation for DNA pools was very low compared to the sampling error (Figure 4). Over nine populations with replicates (excluding Pol3), we observed only four situations among 23,412 loci where two different alleles were fixed in the two replicates (Figure 4). Loci for which an allele was fixed in one replicate was either fixed or displayed a high frequency (above 88%) for the same allele in the other replicate in 98% of cases. Moreover, we estimated pairwise roger’s genetic distance (MRD) based on 23,412 SNPs between the two independent pools from the same landraces. Excluding Population Pol3 (MRD = 0.208), this distance ranged from 0.087 to 0,120 (Table S3). These values provide a reference to decide whether two populations can be considered different or not.

**Table 3:** Sampling error estimated by numerical calculation for one or two biological replicates with independent sampling of 15 or 100 individuals within landraces. Lower and upper bounds indicate the 95% confidence interval for the allelic frequency in the population, based on the binomial probability of the frequency estimated with the corresponding sample size.

| Allelic Frequency | 15 individuals |  |  |  |  |  | 100 individuals |  |  |  |  |  |
| --- | --- | --- | --- | --- | --- | --- | --- | --- | --- | --- | --- | --- |
|  | One biological replicate |  |  | Two biological replicates |  |  | One biological replicate |  |  | Two biological replicates |  |  |
|  | # Alleles | Lower bound | Upper bound | # Alleles | Lower bound | Upper bound | # Alleles | Lower bound | Upper bound | # Alleles | Lower bound | Upper bound |
| **0** | 0 | 0 | 0.116 | 0 | 0 | 0.06 | 0 | 0 | 0.018 | 0 | 0 | 0.009 |
| **0.03** | 1 | 0.001 | 0.172 | 2 | 0.004 | 0.115 | 6 | 0.011 | 0.064 | 13 | 0.017 | 0.055 |
| **0.1** | 3 | 0.021 | 0.265 | 6 | 0.038 | 0.205 | 20 | 0.062 | 0.15 | 40 | 0.072 | 0.134 |
| **0.2** | 6 | 0.077 | 0.386 | 12 | 0.108 | 0.323 | 40 | 0.147 | 0.262 | 80 | 0.162 | 0.243 |
| **0.3** | 9 | 0.147 | 0.494 | 18 | 0.189 | 0.432 | 60 | 0.237 | 0.369 | 120 | 0.256 | 0.348 |
| **0.4** | 12 | 0.227 | 0.594 | 24 | 0.276 | 0.535 | 80 | 0.332 | 0.472 | 160 | 0.352 | 0.45 |
| **0.5** | 15 | 0.313 | 0.687 | 30 | 0.368 | 0.632 | 100 | 0.429 | 0.571 | 200 | 0.45 | 0.55 |
| **0.6** | 18 | 0.406 | 0.773 | 36 | 0.465 | 0.724 | 120 | 0.529 | 0.669 | 240 | 0.55 | 0.648 |
| **0.7** | 21 | 0.506 | 0.853 | 42 | 0.568 | 0.812 | 140 | 0.631 | 0.763 | 280 | 0.653 | 0.745 |
| **0.8** | 24 | 0.614 | 0.923 | 48 | 0.677 | 0.892 | 160 | 0.738 | 0.853 | 320 | 0.757 | 0.838 |
| **0.9** | 27 | 0.735 | 0.979 | 54 | 0.795 | 0.962 | 180 | 0.85 | 0.938 | 360 | 0.866 | 0.928 |
| **1** | 30 | 0.884 | 1 | 60 | 0.94 | 1 | 200 | 0.982 | 1 | 400 | 0.991 | 1 |

**Figure 3:**
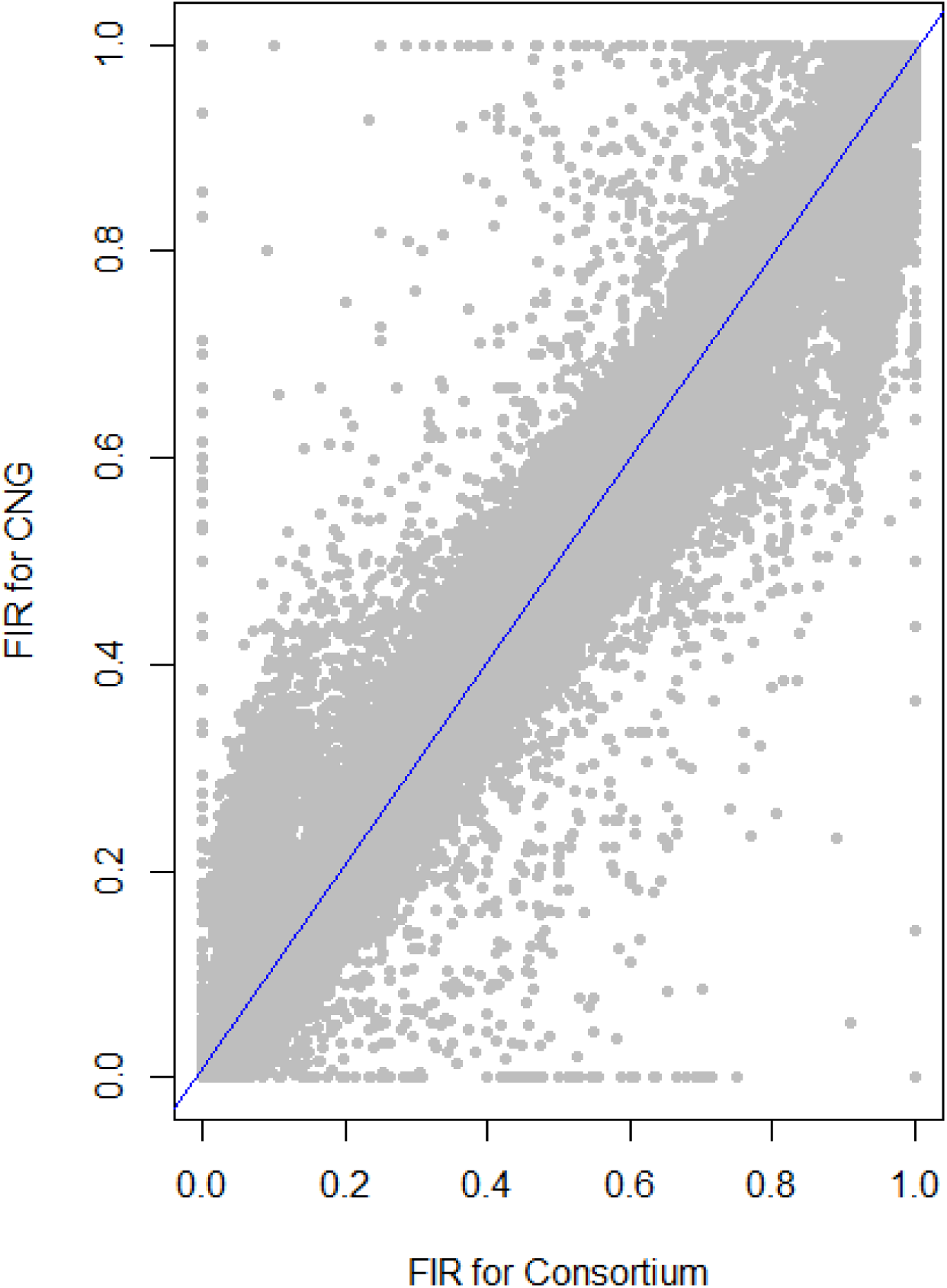
Relationship between fluorescent intensity ratio of European Flint controlled pools genotyped in two different laboratories: CNG and Consortium. Each dot represents the combination of one out 9 controlled pools and one out of 23,412 PZE SNPs. Coefficient of determination (r^2^) between FIR of two laboratories is 0.987.

**Figure 4:**
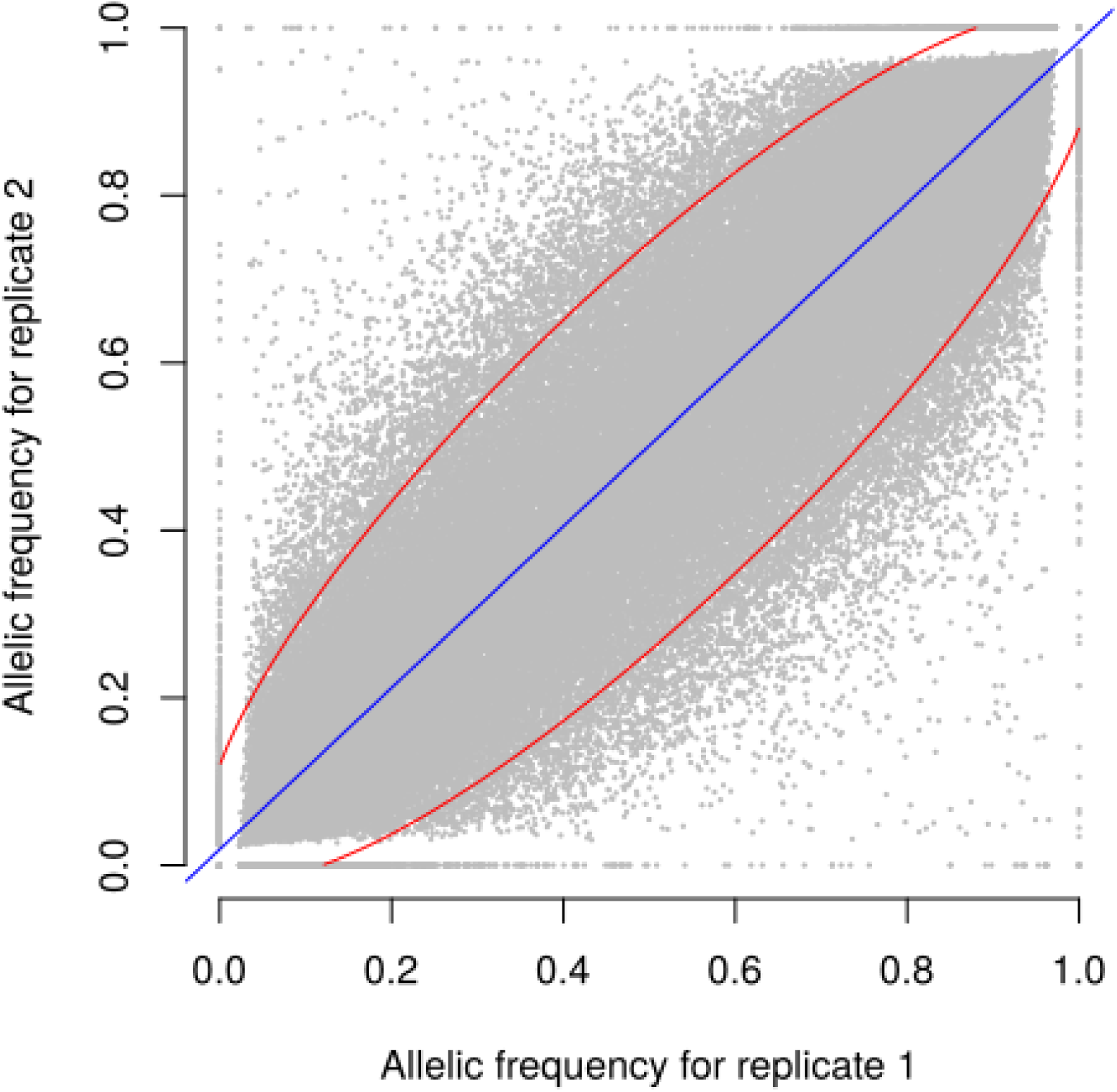
Relationship between allele frequencies predicted for two biological replicates of 9 landraces over 23,412 selected SNPs. Each dot represents one landrace and one SNP, with allele frequency of replicates 1 and 2 on X and Y axes, respectively. Blue line indicates linear regression. 94.5% of points are included in the red ellipse that represents the 95% confidence limit accounting for the effect of sampling alone. r^2^ between replicates is 0.93

### Effect of SNP number and wd on the relationship of genetic distance estimated with SNP and SSR

Finally, we evaluated the possible ascertainment bias due to SNP selection with our filtering based on *wd* criterion. MRD obtained with 17 SSR markers (MRD_SSR_) and MRD based on different set of SNP markers (MRD_SNP_) were highly correlated (Figure 5), indicating a low ascertainment bias. The selection of SNPs based on *wd* quality criterion strongly increased the coefficient of determination (r^2^) between MRD_SNP_ and MRD_SSR_, from 0.587 to 0.639 (Figure S6). We attempted to define the minimal SNP number required to correctly describe the relationship between maize landraces. While increasing the number of SNPs from 500 to 2500 slightly increased r^2^ between MRD_SNP_ and MRD_SSR_ from 0.606 to 0.638 (Figure S6 D,E,F), we observed no further increase beyond 2500 SNPs (Figure S6 A, B, C) suggesting that 2,500 SNPs are enough to obtain a correct picture of landrace relationships.

**Figure 5:**
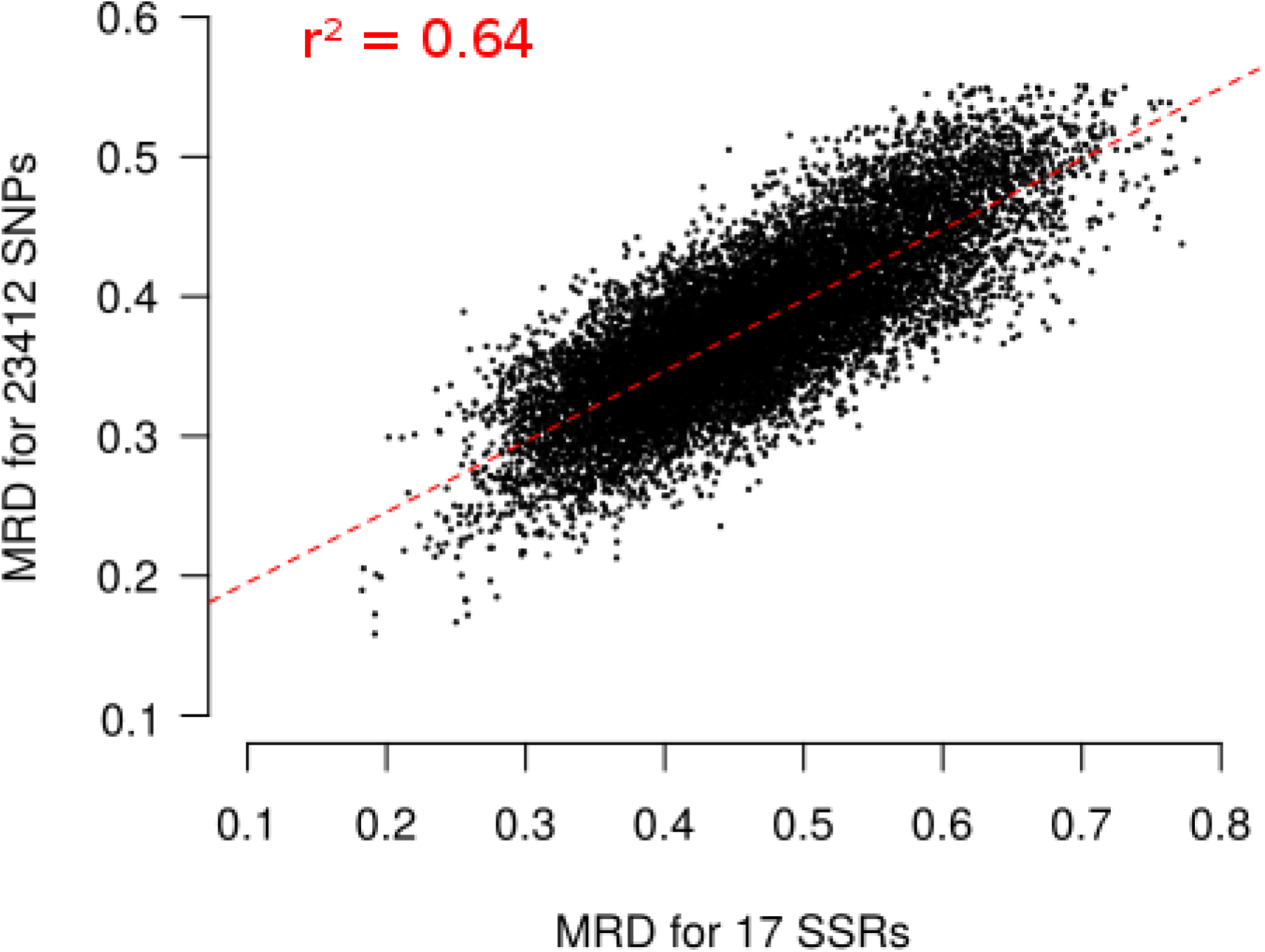
Relationship between Modified Roger’s Distances (MRD) obtained with 17 SSRs and 23,412 SNPs for 156 landraces. Each dot represents one pair of landraces. Red dotted lines represents linear regression between MRD_SSR_ and MRD_SNP_. Coefficient of determination (r^2^) is reported on the plot.

## DISCUSSION

A molecular approach for diversity analysis of landraces needs to answer several criteria (i) an accurate estimation of allelic frequency in each population, (ii) a robust and reproducible measurement of allelic frequency across laboratories in order to facilitate comparison of genetic diversity of accessions across genebanks, (iii) a reliable estimate of genetic distance between landraces with no or little ascertainment bias (iv) an affordable, high-throughput and labor efficient method, due to both strong financial and human constraints in plant genebanks. Four main sources of errors affect the accuracy of allelic frequency estimation of a locus in a population using a DNA pooling approach: (i) the sampling of individuals (so called “sampling” errors), (ii) the procedure to mix DNA from individuals (so called “DNA mixing” errors) (iii) the imprecision of quantitative measurement used by the model for the prediction (so called “experimental” errors) and (iv) the predictive equation used to predict allelic frequency in a population (so called “approximation” errors).

### A two-step model to protect against erroneous detection of polymorphism and predict accurately allelic frequencies in DNA bulk

Approximation errors due to predictive equation depend on (i) the model used to predict allelic frequencies and (ii) the set of individuals and SNPs used to calibrate the predictive equation. In this study, we used a two-step modeling using inbred lines and controlled pools as sets of calibration to test for polymorphism and then predict allelic frequency for polymorphic markers. Detection of allele fixation in a population is an important issue for deciphering and managing genetic diversity within plant and animal germplasm. We used two Student tests based on fluorescent intensity ratio (FIR) distribution of lines homozygous for allele A and B to determine polymorphism of a SNP in a given landrace (Figure 1). In this first step, we preferred a method based on FIR distribution rather than the clustering approach implemented in genome studio because it is possible to control the type I risk of false allele detection (at 5% in our study). Using this two-step approach reduces strongly the erroneous detection of polymorphisms in a population compared to previous methods: MAE for fixed locus <0.1% in our approach (Table S2) vs ∼2-3% using PPC method (Brohede 2005) or ∼2-8% using different k correction from (Peiris *et al*. 2011). This is not surprising as previous methods focused on an accurate estimation of the difference in allele frequencies between DNA bulks of individuals contrasted for a quantitative trait of interest (Sham *et al*. 2002; Craig *et al*. 2005; Kirov *et al*. 2006; Teumer *et al*. 2013) and did not focus specifically on protecting again false detection of alleles.

For loci that were detected as polymorphic, we predicted allelic frequencies from FIR in landrace DNA pools by using a unique logistic regression for 23,412 SNPs passing *wd* quality criterion. The relationship between FIR and allelic frequency was modelled using a quasi-logistic regression for different reasons. First, the logistic function ensures that the predicted frequencies take value in (0,1), a property that is not satisfied by polynomial regression (PPC) or tan transformation (Brohede 2005; Teumer *et al*. 2013). Second, one could observe that the relationship between the fluorescent intensity ratio and allelic frequencies within controlled pools was not linear (Figure S2).

This two-step approach led to a low global error rate in allelic frequency prediction (MAE = 3% for polymorphic and monomorphic loci considered jointly; Figure 2, Table S2). It is comparable to the most accurate previous pooling DNA methods for SNP array that used a specific model for each SNP: (i) MAE ranging from 3 to 8 % (Peiris et al., 2011) or 5-8% (Brohede et al., 2005) depending of k-correction applied (ii) MAE ∼ 3% for PPC correction (Brohede et al., 2005; Teumer et al., 2013) (iii) MAE ∼ 1% for tan-correction (Teumer et al., 2013). Several factors can explain this relative good global accuracy of our approach. First, almost half of the loci were fixed on average in each landrace, which contributed positively to global accuracy since our method over-performed previous methods for fixed locus (see above). Second, *wd* quality criterion removed SNPs for which allelic frequencies were poorly predicted using FIR. We observed indeed that increasing the threshold for *wd* quality criterion led to a global increase in accuracy at both steps (Figure S1). While 90% of SNPs have a MAE<2% for wd criterion >10, only 50% of SNPs have a MAE<2% for wd criterion <10. Taking into account differential hybridization by using a specific logistic regression for each SNP could be a promising way to further improve the accuracy of allelic frequencies prediction, notably for balanced allelic frequencies (Brohede et al., 2005, Peiris et al., 2011, Teumer et al., 2013). To limit possible ascertainment bias and errors in allelic frequency estimation, it requires however to genotype additional series of controlled pools for SNPs for which current controlled pools were monomorphic or have a limited range of allelic frequency (Figure S3 and S4).

To estimate the parameters of the logistic regression, we used two series of controlled pools rather than heterozygous individuals for both technical and practical reasons. Controlled pools cover more homogenously the frequency variation range than heterozygous and homozygous individuals only, which therefore limits the risk of inaccurate estimation of logistic model parameters. Different studies showed that accuracy of allelic frequency estimation strongly depends on accuracy of FIR estimation for heterozygous individual and therefore the number of heterozygous individuals (Le Hellard *et al*. 2002; Simpson 2005; Jawaid and Sham 2009). Between 8 and 16 heterozygous individuals are recommended to correctly estimate FIR mean for heterozygous individuals, depending on FIR variance (Le Hellard *et al*. 2002). In maize, we can obtain heterozygote genotypes either by crossing inbred lines to produce F1 hybrids, by planting seeds from maize landraces, or by using residual heterozygosity of inbred lines. Using residual heterozygosity to calibrate model is not possible since half SNPs show no heterozygous genotype in the 327 inbred lines of our study. Obtaining at least 16 heterozygous individuals for each SNP therefore requires to genotype a few dozens of F1 hybrids or individuals from landraces considering that expected heterozygosity in a landrace is comprised between 3 and 28% (Arca et al., in prep). This represents additional costs since maize researchers and breeders genotyped preferentially inbred lines to access directly haplotypes without phasing and because genotypes of F1 hybrids can be deduced of that of their parental inbred lines. Beyond allogamous species as maize, genotyping heterozygous individuals could be time demanding and very costly in autogamous cultivated plant species for which genotyped individuals are mostly homozygotes (wheat, tomato, rapeseed). On the contrary, one can easily produce controlled pools whatever the reproductive system, either by mixing DNA or equal mass of plant materials, which allows producing a wide range of allelic frequencies.

### Effect of DNA mixing procedure on accuracy allelic frequency estimation

There are two main errors coming from DNA mixing procedure: (i) the “sampling error” that is directly connected to the number of individuals sampled in each population (Table 3), and (ii) the “bulking error” associated with the laboratory procedure to mix equal DNA amounts of sampled individuals.

We evaluated sampling and bulking errors by comparing 10 independent biological replicates from 10 different landraces obtained by independently sampling and mixing equal leaf areas of young leaves of 15 individuals. Allelic frequencies estimated for both biological replicates from a same landrace were highly correlated. Excluding Pol3, 94.5% of difference of allelic frequencies between replicates was of included within 95% confidence limits originated from sampling effect only Figure 4). This suggests that the “bulking error” is low compared to the “sampling error”. Consistently, Dubreuil et al., (1999) observed a low “bulking error” for RFLP markers using the same DNA pooling method, with a coefficient of determination of 0.99 between allelic frequencies based on individual genotyping of plants and those predicted using DNA bulks. Several studies also showed that the effect of bulking errors on allelic frequencies measured by comparing DNA pool and individual genotyping of plant of this DNA pool is very low compared with other sources of errors (Le Hellard *et al*. 2002; Jawaid and Sham 2009). Additionally, the mixing procedure starting from leaf samples strongly reduced the number of DNA extractions for each DNA bulk as compared to first extracting DNA from each individual, and then mixing by pipetting each DNA samples to obtain an equimolar DNA mix (“post-extraction” approach). Since the cost of DNA extraction becomes non-negligible when the number of individual increases, mixing plant material based on their mass before extraction is highly relevant to save time and money. This can be done without losing accuracy as shown in this study for SNP array and previously for RFLP by Dubreuil et al., (1999).

We highlighted the critical importance of the number of individuals sampled per landrace on allelic frequency estimation (Table 3). By using DNA pooling, accuracy can be gained with very little additional cost by increasing number of sampled individuals. Whereas a high accuracy of allelic frequency estimation within landraces is required to scan genome for selective sweeps, it is less important to estimate global genetic distance, due to the large number of SNPs analyzed. Sampling fifteen plants per population (30 gametes) appears convenient to obtain an accurate estimation of frequencies in a population and analyze genetic diversity (Reyes-Valdés *et al*. 2013).

### A low ascertainment bias to estimate genetic distance between landraces

There are two possible sources of ascertainment bias using a DNA pooling approach on a SNP array. The first one relates to the design of array because the set of lines to discover SNPs may not well represent genetic diversity and a threshold in allelic frequency was possibly applied to select SNPs. The second one relates to the selection of a subset of SNPs from the array regarding the genetic diversity of samples in calibration set used to predict allelic frequencies.

To avoid risk of ascertainment bias due to selection of markers genotyped by the array, the logistic regression model was adjusted on 1,000 SNPs with the largest allelic frequency range rather than for each of the 23,412 PZE SNPs individually. Using a specific model for each SNP would indeed conduct to discard markers monomorphic in controlled pools and therefore select only markers polymorphic between parents of controlled pool. Note that the same issue would be raised by using heterozygous individuals since 8 to 16 heterozygotes were recommended to adjust a logistic regression. Using heterozygous individuals and SNP specific equations could lead to systematically counter-select SNPs with low diversity. It could also lead to systematically remove SNPs that are differentially fixed between isolated genetic groups, because no or very few heterozygote individuals are available.

We also evaluated ascertainment bias by comparing Modified Roger’s Distance (MRD) between the 156 landraces obtained using SNPs (MRDSNP) and SSRs (MRDSSR) (Camus-Kulandaivelu 2006; Mir *et al*. 2013), which display no or limited ascertainment bias. MRDSNP was highly correlated with MRDSSR (r2 = 0.64; Figure 5). This correlation is high considering that SSR and SNP markers evolve very differently (mutation rate higher for SSRs than SNPs, multiallelic vs biallelic), that the number of SSR markers used to estimate genetic distance is low and that errors in allelic frequency prediction occur for both SNPs and SSRs. For comparison, correlation was lower than between Identity By State estimated with 94 SSRs and 30k SNPs in a diversity panel of 337 inbred lines (r2 = 0.41), although very few genotyping errors are expected in inbred lines (Bouchet *et al*. 2013). Using the *wd* criterion significantly increased the correlation between MRDSNP and MRDSSR markers for 156 landraces (Figure S5). It suggests that the *wd* criterion removes SNP markers that blurred the relationships between landraces. We can therefore define a subset of 23,412 SNPs to analyze global genetic diversity in landraces. This is in agreement with previous studies in inbred lines showing that PZE SNPs are suitable to analyze the genetic diversity in inbred lines (Inghelandt *et al*. 2011; Ganal *et al*. 2011; Bouchet *et al*. 2013; Frascaroli *et al*. 2013). These studies showed that diversity analysis based on PZE SNPs give consistent results with previous studies based on SSR markers (Inghelandt *et al*. 2011; Bouchet *et al*. 2013; Frascaroli *et al*. 2013).

The DNA pooled-sampling approach therefore provides a reliable picture of the genetic relatedness among populations that display a large range of genetic divergence and opens a way to explore genome-wide diversity along the genome.

### An affordable, high-throughput, labor-efficient and robust method compared to SSR / RFLP markers and sequencing approaches

Using SNP arrays instead of SSR/RFLP marker systems or sequencing approaches has several advantages. First, SNP genotyping using arrays is very affordable compared to SSR/RFLP or resequencing approaches because it is highly automatable, high-throughput, labor-efficient and cost effective (currently 30-80€ / individual depending of array). Obtaining accurate estimations of allelic frequencies using a whole genome sequencing (WGS) approach requires high depth and coverage for each individual because of the need of counting reads (Schlötterer *et al*. 2014). To estimate allelic frequency in DNA bulks, WGS remains costly compared to SNP arrays for large and complex genomes of plant species as maize. Different sequencing approaches based either on restriction enzyme or sequence capture make it possible to target some genomic regions and multiplex individuals, reducing the cost of library preparation and sequencing while increasing the depth for the selected regions (Glaubitz *et al*. 2014). However, these sequencing approaches remain more expensive than SNP arrays and require laboratory equipment to prepare DNA libraries and strong bioinformatics skills to analyze sequencing data. These skills are not always available in all genebanks. With the maize 50K array, FIR measurement used to predict allelic frequencies were highly reproducible both across laboratories and batches (r2 = 0.987; Figure 3). We can therefore consistently predict allelic frequencies using 50K array in new DNA pools genotyped in other laboratories, by applying the same parameters of presence /absence test and logistic regression as in this study. This will greatly facilitate the comparison of accessions across collections and laboratories. This is a strong advantage over SSRs for which a strong laboratory effect has been observed for the definition of alleles, leading to difficulties for comparing genetic diversity across seedbanks and laboratories (Mir *et al*. 2013). Similarly, one can expect some laboratory effect for sequencing approaches due to preparation of library and bioinformatics analysis. However, there is some disadvantage to use SNP arrays instead of SSR markers or sequencing approach. First, SNP marker are bi-allelic whereas SSRs are multi-allelic. At a constant number of markers, using SNPs rather than SSRs therefore leads to less discriminative power (Laval *et al*. 2002; Hamblin *et al*. 2007). This disadvantage is largely compensated by the higher number of SNPs and the fact that SNPs are more frequent and more regularly spread along the genome than SSR/RFLP, allowing genome wide diversity analyses. Second, contrary to SSR / RFLP markers and sequencing approach, SNP array does not allow one to discover new polymorphisms, which may lead to ascertainment bias for diversity analysis of new genetic groups (Nielsen 2004; Clark *et al*. 2005; Hamblin *et al*. 2007; Inghelandt *et al*. 2011; Frascaroli *et al*. 2013). Comparison with SSRs results showed that PZE SNPs provide reliable genetic distances between landraces, suggesting a low ascertainment bias for a global portrayal of genetic diversity (see above). Sequencing techniques may be interesting in a second step to identify, among preselected accessions, those which show an enrichment in new alleles.

The number of SNPs affects the estimates of relationship between landraces and population structure (Moragues *et al*. 2010). In our study, the correlation coefficient between MRDSNP and MRDSSR increased with increasing number of SNPs and reached a plateau for 2,500 SNPs (Figure S6). This suggests that increasing the number of SNPs above 2,500 does not provide further improvement in precision to estimate relationships between landraces as compared to 17 SSRs. Our approach could therefore be made further cost efficient by selecting less loci for studying global genetic relationships and genetic diversity. For maize, a customizable 15K Illumina genotyping array has been developed that includes 3,000 PZE SNPs selected for studying essential derivation (Rousselle *et al*. 2015) and 12,000 others selected for genetic applications such as genomic selection. Alternatively, the same approach could be applied to other genotyping arrays with higher density as the 600K Affymetrix Axiom Array (Unterseer et al., 2013) to gain precision in detection of selective footprints.

## CONCLUSION

The DNA pooling approach we propose overcomes specific issues for genetic diversity analysis and plant germplasm management purposes that were not or partially addressed by previous methods which were mostly focused on QTL analysis and genome wide association studies (Hoogendoorn *et al*. 2000; Craig *et al*. 2005; Brohede 2005; Teumer *et al*. 2013). As proof of concept, we used the DNA pooling approach to estimate allelic frequencies in maize landraces in order to identify original maize landraces in germplasm for pre-breeding purposes and selective footprints between geographic and/or admixture groups of landraces cultivated in contrasted agro-climatic conditions (Arca et al., in prep). Our approach could be very interesting for studying plant germplasm since time, money and molecular skills can be limiting factors to study and compare large collections of landraces maintained in seedbank (Mir *et al*. 2013). Applications could be expanded to QTL identification in pools (Gallais *et al*. 2007), detecting signatures of selection in multi-generation experiments, or detection of illegitimate seed-lots during multiplication in genebanks. The DNA pooling approach could be easily applied to decipher organization of genetic diversity in other plant germplasm since Infinium Illumina HD array have been developed for several cultivated plant species, including soybean, grapevine, potato, sweet cherry, tomato, sunflower, wheat, oat, brassica crops but also animal species.

## MATERIALS AND METHODS

### Plant material

#### Landraces

A total of 156 landrace populations (Table S4) were sampled among a panel of 413 landraces capturing a large proportion of European and American diversity and analyzed in previous studies using RFLP (Dubreuil and Charcosset 1998; Rebourg *et al*. 1999, 2001, 2003; Gauthier *et al*. 2002) and SSR markers (Camus-Kulandaivelu 2006; Dubreuil *et al*. 2006; Mir *et al*. 2013).

Each population were represented by a bulk of DNA from 15 individual plants, mixed in equal amounts as described in Reif *et al*. (2005) and Dubreuil *et al*. (2006). In order to analyze the effect of individual sampling on allelic frequency estimation (see below), ten populations were represented by two DNA bulks of 15 plants sampled independently (Table 3).

#### Controlled DNA Pools

To calibrate a prediction model for SNP allelic frequencies in populations, we considered two series of nine controlled pools derived from the mixing of two sets of three parental inbred lines: EP1 – F2 – LO3 (European Flint inbred lines) and NYS302– EA1433 – M37W (Tropical inbred lines).

For each set of three parental lines, we prepared nine controlled pools by varying the proportion of each line in the mix (Table 1), measured as the number of leaf disks with equal size according to Dubreuil et al., (1999). The proportion of lines 2 and 3 (EA1433 and M37W or F2 and LO3) varies similarly whereas line 1 (EP1 or NYS302) varies inversely. The genotype of the inbred lines and the proportion of each inbred line in each pool give the expected allelic frequencies as shown in Table 1. Combination of genotypes in parental lines can conduct either to monomorphic or polymorphic controlled pools if the genotypes of 3 parental lines are the same or not, respectively. If we exclude monomorphic controlled pools and heterozygote SNPs in parental lines, these different combinations conduct to four different polymorphic configurations in the 9 controlled pools, corresponding to four ranges of allelic frequencies: 1-33% (R1), 33-50% (R2), 51-67% (R3), 67-99% (R4), (Table 1). Combination of R1 and R4 configurations in two series of controlled pools displayed the largest allelic frequencies range (1% to 99%) while combination of R2 and R3 displayed a more reduced allelic frequency range (33% to 67%).

#### Inbred lines

To test for allele fixation within landraces, we used a panel of 333 inbred maize lines representing the worldwide diversity well characterized in previous studies (Camus-Kulandaivelu 2006; Bouchet *et al*. 2013) (Table S5). This panel includes the six inbred lines used to build two series of controlled pools.

### Genotyping

We used the 50K Illumina Infinium HD array (Ganal *et al*. 2011) to genotype (i) 166 DNA bulks representing 156 landraces (ii) 18 DNA bulks representing 2 series of controlled DNA pools (iii) 333 inbred lines. 50K genotyping was performed according to the manufacturer’s instructions using the MaizeSNP50 array (IlluminaInc, San Diego, CA). The genotype results were produced with GenomeStudio Genotyping Module software (v2010.2, IlluminaInc) using the cluster file MaizeSNP50_B.egt available from Illumina. The array contains 49,585 SNPs passing quality criteria defined in (Ganal *et al*. 2011).

We also used 17 SSRs genotyping data from 145 and 11 landraces analyzed by Camus et al. (2006) and Mir et al. (2013), respectively.

#### Measurement variable: fluorescence intensities ratio

The MaizeSNP50 array has been developed into allele-specific single base extension using two colors labeling with the Cy3 and Cy5 fluorescent dyes. The fluorescent signal on each spot is digitized using GenomeStudio software. Data consist of two normalized intensity values (x, y) for each SNP, with one intensity for each of the fluorescent dyes associated with the two alleles of the SNP. The alleles measured by the x intensity value (Cy5 dye) are arbitrary, with respect to haplotypes, are called the A alleles, whereas the alleles measured by the y intensity value (Cy3 dye) are called the B alleles.

We assumed that the strength of the fluorescent signal of each spot is representative of the amount of labeled probe associated with that spot. The amount of labeled probes at each spot relies upon the frequency of the corresponding alleles of PCR product immobilized on it. Based on this assumption, the fluorescent intensity ratio (FIR) of each spot (y/(x+y)) can be employed to estimate the allele frequency of DNA bulk immobilized on it.

To test the reproducibility of the measurement the controlled pool of European lines was genotyped twice in two platforms, at CNG Genotyping National Center, Evry 91, France, and at Trait Genetics.

#### SNP filtering and quality control

For the purpose of this study, we used only the subset of 32,788 markers contributed by the Panzea project (http://www.panzea.org/), so called PZE SNPs, developed on the basis of US NAM founders (Zhao 2006). These SNPs represent a comprehensive sample of the maize germplasm and are therefore suitable for diversity analysis (Ganal *et al*. 2011).

The following equation (1) was then used to create a rank score (weighted deviation, wd) for each SNP in order to identify and remove those of poor quality,

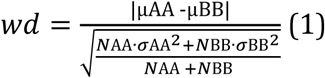

where µ_*AA*_ and s_AA_ and µ_*BB*_ and s_BB_ are the mean and the standard deviation for the fluorescence intensity ratios of AA and BB genotypes for the 327 inbred lines panel and N_AA_ and N_*BB*_ is the number of inbred lines with genotype AA or genotype BB respectively. To avoid selection bias, loci which were monomorphic within the reference inbred lines population were selected using the *wd* equations (1), assuming µ_*AA*_*=*0 and s_AA_*=*0 for monomorphic BB SNPs or assuming µ_*BB*_*=1* and s_BB_*=*0 for monomorphic AA SNPs.

This criterion removes from analysis those SNPs for which distributions of fluorescence signal ratios for AA and BB genotypes of 327 inbred lines panel overlap or have large variances. To analyze genetic diversity, we first selected 23,656 with *wd* above 50 among 32,788 PZE SNPs. This threshold removed SNPs displaying high error rate in allelic frequency prediction (Figure S1). In addition, we removed 244 SNPs that were heterozygous in one of parental lines of controlled pools and that displayed high error rate in allelic frequency prediction (data not shown).

#### Alleles detection and allele frequency estimation

Allele frequency estimation within DNA pools was implemented as a two-step process. We first determined the fixation of alleles A and/or B by comparing the fluorescent ratio of DNA pools at a given SNP locus with the distribution of the fluorescent signal of inbred lines (see above) which have AA or BB genotypes at the same locus. We assumed Gaussian distributions for the fluorescent intensities and tested for fixation using a Student’s t-tests with a 5% type I nominal level.

In second step, for each SNP for which alleles A and B were both declared present, the allelic frequency fB of allele B was inferred using the following generalized linear model:

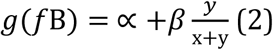

where *x* and *y* are the fluorescent intensities at SNP for alleles A and B respectively, α and β are the parameters of a logistic curve, calibrated on fluorescent ratio data from controlled pools for 1000 SNPs and ε_*i*_ is a noise term. As allele B frequency is a binomial variable, GLM was set with a logit link function (R, version 3.0.3).

The calibration sample of 1,000 SNPs consists in 250 randomly selected SNPs for each possible configuration (R1, R2, R3, R4 defined in Table 1). It was preferred to a calibration sample of all SNPs or to a specific prediction curve for each SNPs, in order to have a homogeneous distribution of observations into each class of expected frequency. Calibrating the model for each SNP would lead to high error in allelic frequency prediction, notably for monomorphic controlled pools as exemplified by Figure S3 and S4. Calibrating model for all SNPs would give strong weight to fixed allele in calibration due to large number of monomorphic controlled pools that are homozygous either for allele A or B.

### Accuracy of allelic frequency estimation

We assessed the accuracy of allele frequency estimates from pooled DNA samples by calculating the absolute difference between allelic frequencies of the B allele predicted by our two-step model and those expected for controlled pools from the genotype of their six parental lines. We obtained expected allelic frequencies for two series of controlled pools by weighting the allelic frequency of each parental line (0 or 1) by their relative mass in the mix (Table 1). We obtained genotypes of inbred lines from clustering by genome studio. This absolute difference was averaged over SNPs and samples in order to obtain mean absolute error (MAE).

We first evaluated the mean absolute error for 23,412 SNPs in the two series of controlled pools (Table S2, Figure 2). In order to estimate the effect of the calibration set of individuals and SNPs on the accuracy of allelic frequency prediction, we applied two cross-validation approaches on the 1000 SNPs and the two series of controlled pools and six parental inbred lines (24 samples) used to calibrate parameters of the common logistic regression. In order to evaluate the effect of SNP calibration set (Table S1), we repeated five time a K-fold approach in which 1000 SNPs were split randomly in a training set of 800 SNPs on which we calibrated our two-step model and a validation set of 200 SNPs on which we predicted allelic frequency using this model in same two series controlled pools and estimated MAE. In order to evaluate the effect calibration samples (Table 2), we repeated 1000 times a K-fold approach on 1000 SNPs in which 1, 3, 5, 8, 10, 15 samples among 18 from controlled pools were randomly removed from the calibration set. We used the remaining samples to estimate parameters of the logistic regression, and then predicted allelic frequencies using this predictive equation in these K removed samples (Table 2).

To estimate sampling error (Table 3), we estimated the 95% confidence interval of the allelic frequency in the population considering various observed allelic frequency obtained by sampling either 15, 30, 100 or 200 individuals from this population. To obtain the lower and upper bound of the 95% confidence interval for allelic frequency in the population, we considered the binomial probability to obtain various number of allele B in 15, 30, 100, 200 individuals (estimated allelic frequencies) from a population (true allelic frequencies) by using binom.confint function implemented in R package “binom”. We used the following parameters: binom.confint(x = number of alleles observed, n = 2*number of individuals, conf.level=95%, methods = exact) with x = number of successes and n = number of trial in the binomial experiment.

### Comparison of genetic distance between SNP and SSR markers

We calculated the modified Roger’s distance (MRD) (Rogers 1972) based on allelic frequency data between landraces using different sets of markers to analyze the effect of the *wd* criterion (Figure S5) and of the number of markers (Figure S6) on the estimation of relatedness. To analyze the effect of *wd* criterion, we selected four random sets of 2,000 SNPs with different *wd* ranges (0-20, 20-40, 40-60, 60-80) among 32,788 PZE SNPs. To analyze the effect of SNP number, we selected six random sets of SNPs with various number of SNPs (15,000, 10,000, 5000, 2500, 1000, 500) among 23,412 SNPs with *wd* above 50. In order to test if the genetic distance is robust when changing the type and the number of markers, we compared MRD between landraces estimated with different SNP datasets with that estimated with 17 SSR markers (Figure 5, Figure S5 and Figure S6). Missing allele frequencies within accession were replaced by corresponding average frequencies within the whole set of accessions before running this analysis. Allelic frequencies of two samples for replicated landraces were averaged before estimating MRD distance except for Pol3 for which one of two samples was removed (WG0109808-DNAH04).

Coefficient of determination between the distance matrices based on different subsets of SNP (MRD_SNP_) and 17 SSR markers (MRD_SSR_) was determined by using linear regression.

## Acknowledgements

This study was funded by l’Association pour l’étude et l’amélioration du mais (PROmais) in the project “Diversity Zea” and French National Research Agencies in project Investissement d’Avenir Amaizing, (ANR-10-BTBR-01). We acknowledge greatly the French maize Biological Ressources Center, PROmais, and INRAE experimental units of St Martin de Hinx and Mauguio for collecting and maintaining Landraces and Inbred lines collection. We greatly acknowledge the colleagues who initially collected these landraces and André Gallais for having initiated these research programs. We also greatly acknowledge Pierre Dubreuil, Letizia Camus-Kulandaivelu, Cecile Rebourg, Céline Mir, Domenica Maniccaci that conducted previous study on these landraces using DNA pooling approach with SSR and RFLP markers. The Infinium genotyping work was supported by CEA-CNG, by giving the INRAE-EPGV group access to its DNA and cell bank service for DNA quality control and to their Illumina genotyping platform. Thanks respectively to Anne Boland and Marie-Thérèse Bihoreau and their staff. We acknowledge the EPGV group, Dominique Brunel, Marie-Christine Le Paslier, Aurélie Chauveau for the discussion and management of the Illumina genotyping.

## Author’s contribution

S.D.N, A.C and B.G designed and supervised the study and selected the plant material M.A, S.D.N, A.C drafted and corrected the manuscript

D.M, V.C and A.B extracted DNA and managed genotyping of landraces and inbred lines C.B, B.G and A.C collected, maintained landraces, and inbred lines collection

S.D.N, M.A, A.C and T.M-H developed the statistical methods and scripts for predicting allelic frequency from fluorescent data

M.A, B.G and S.D.N analyzed genetic diversity of landraces panel. All authors read and approved the manuscript.

## Data availability

R scripts and fluorescent intensity data of 327 inbred lines and two series of controlled pools used for predicting allelic frequency in DNA bulks of maize landraces by our two-step approaches are available at https://doi.org/10.15454/GANJ7J. Fluorescent Intensity data and allelic frequencies of 20 samples corresponding to 10 duplicated landraces were also available at https://doi.org/10.15454/GANJ7J. Allelic frequencies of new DNA bulks for new maize populations genotyped by maize 50K array could be predicted by using these datasets with R scripts. Note that these datasets and R scripts will become available when the publication would be accepted in a peer review journal.

## Conflicts of interest

No

## Notes

### Competing Interest Statement

The authors have declared no competing interest.

