## Supplementary Figures for "Deciphering the genetic diversity of landraces with high-throughput SNP genotyping of DNA bulks: methodology and application to the maize 50k array"

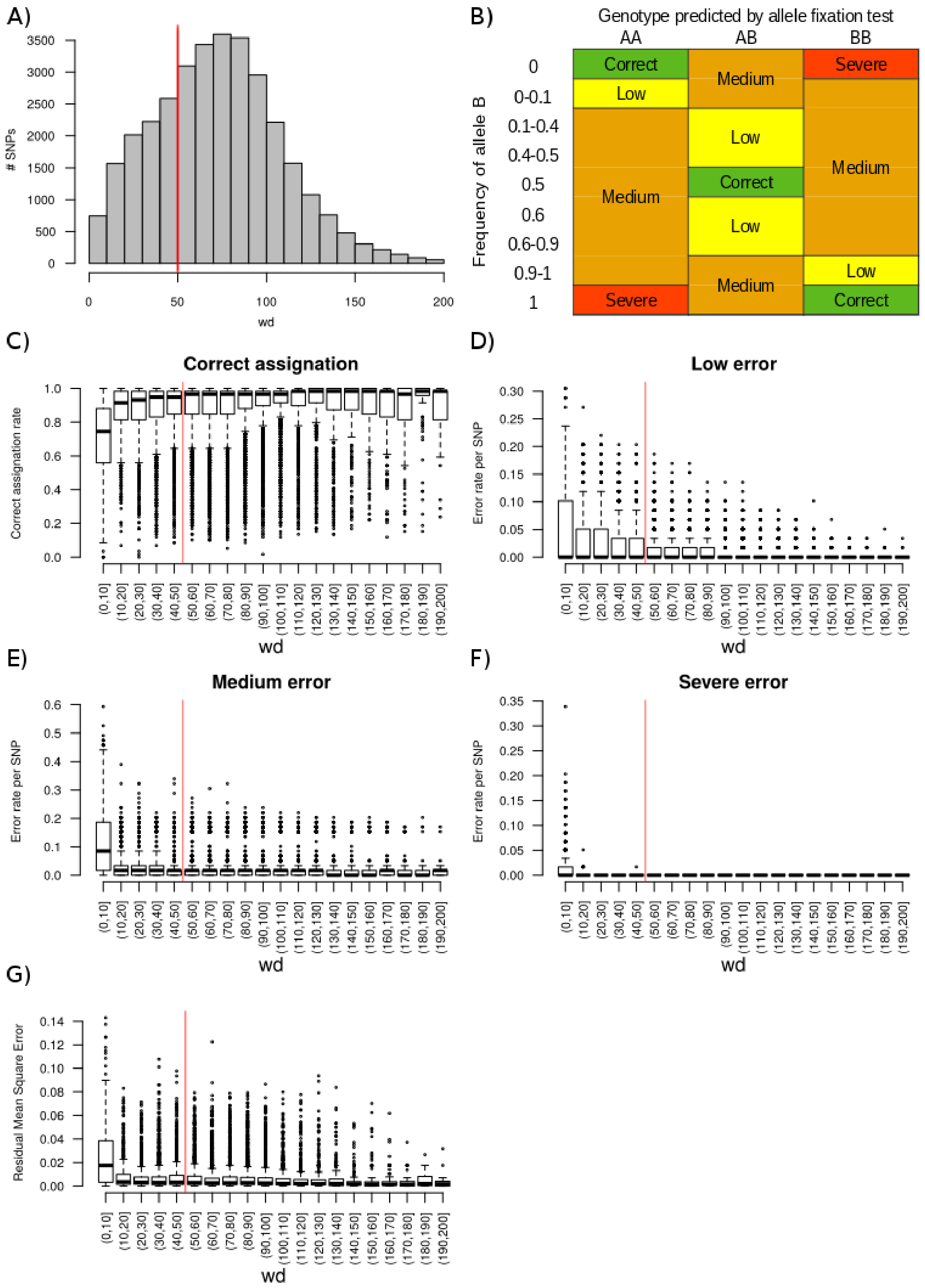


**Figure S1: Effect of weighted deviation (wd) on accuracy of allelic frequency prediction for 32,788 PZE SNPs in controlled pools. A) Distribution of weighted deviation. B) Classification of errors by fixation test, according to the known allelic frequencies in controlled pools. C) Boxplot of correct genotype assignation rate per SNP according to wd. D, E, F) Boxplots of low, moderate and severe genotype assignation error rate per SNP according to wd. G) Boxplot of Residual Mean Square Error (RMSE) obtained by cross-validation, according to wd with a specific logistic regression for a subset of**

**18,670 SNPs. Note that 132 SNPs with wd above 200 were not represented. Red vertical lines represent the wd threshold used to filter SNPs in this study.**

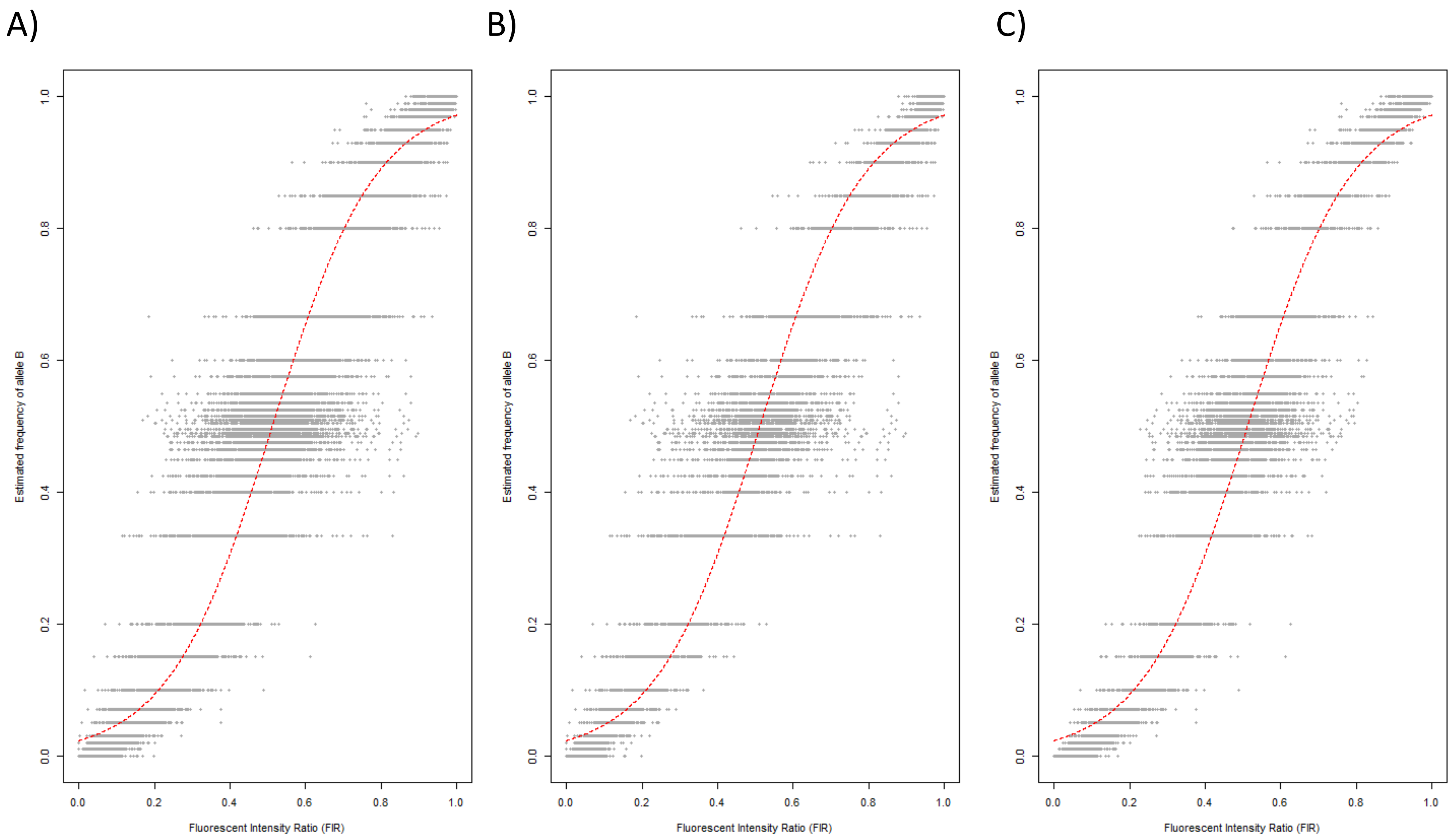


*Figure S2: Relationship between the Fluorescent Intensity Ratio (FIR) and the expected frequency of allele B for 23,412 SNPs in the European and Tropical controlled pools together (A), in the European pools (B) and in the Tropical pools (C).*

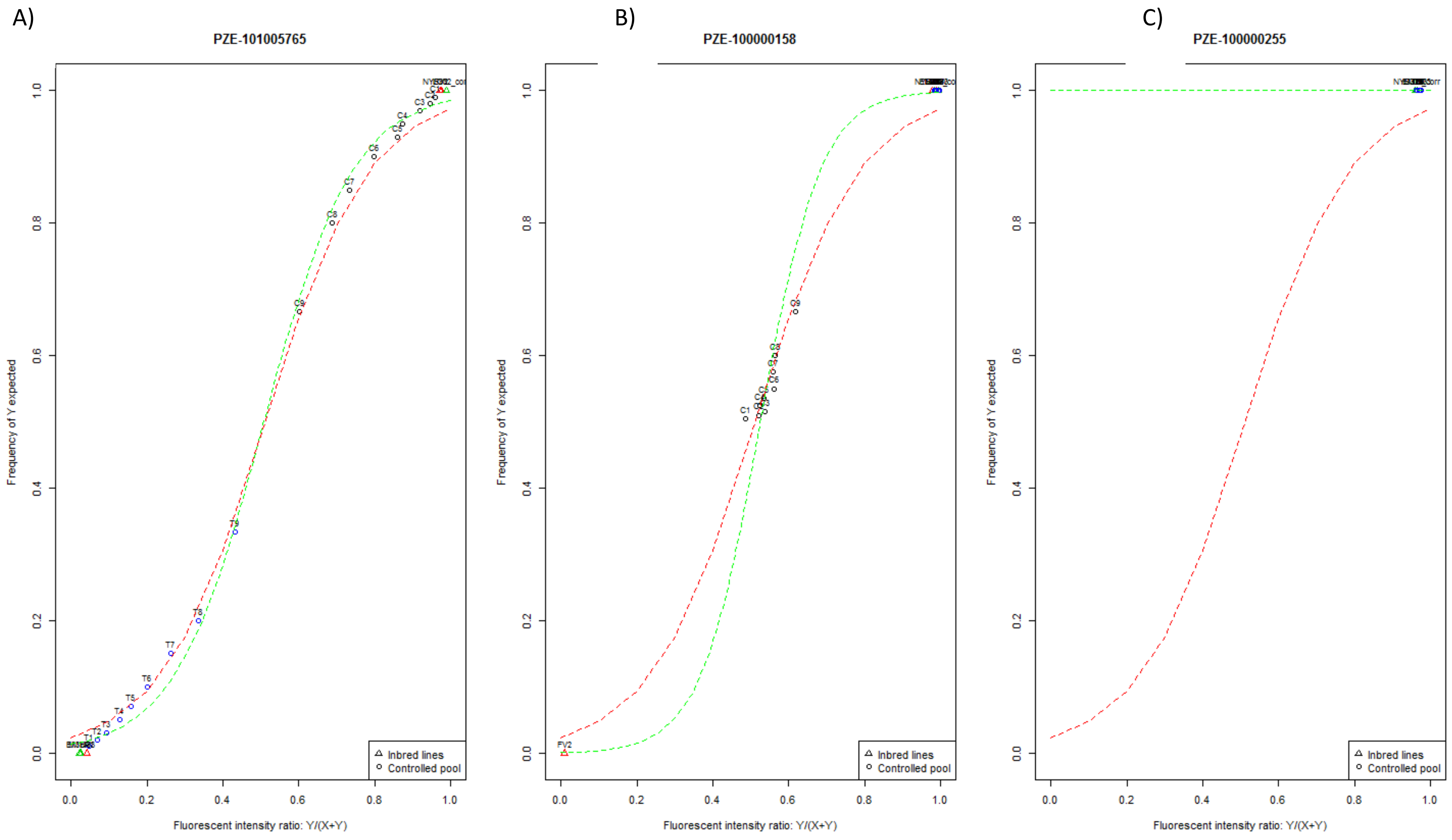


**Figure S3: Common and specific logistic regression curves for three SNPs with different ranges of expected frequency of allele B in controlled pools. Variation in ranges results from the different combination of parental lines genotypes in controlled pools (see Table 1). A) Large allelic frequency range (1%-99%); B) Low allelic frequency range (51%-67%); C) No allelic frequency range in controlled pools corresponds to configurations in which all parental lines have the same genotype. Red and green doted curves represent logistic regression equations calibrated with 1,000 SNPs (common) and with each SNP individually (specific), respectively.**

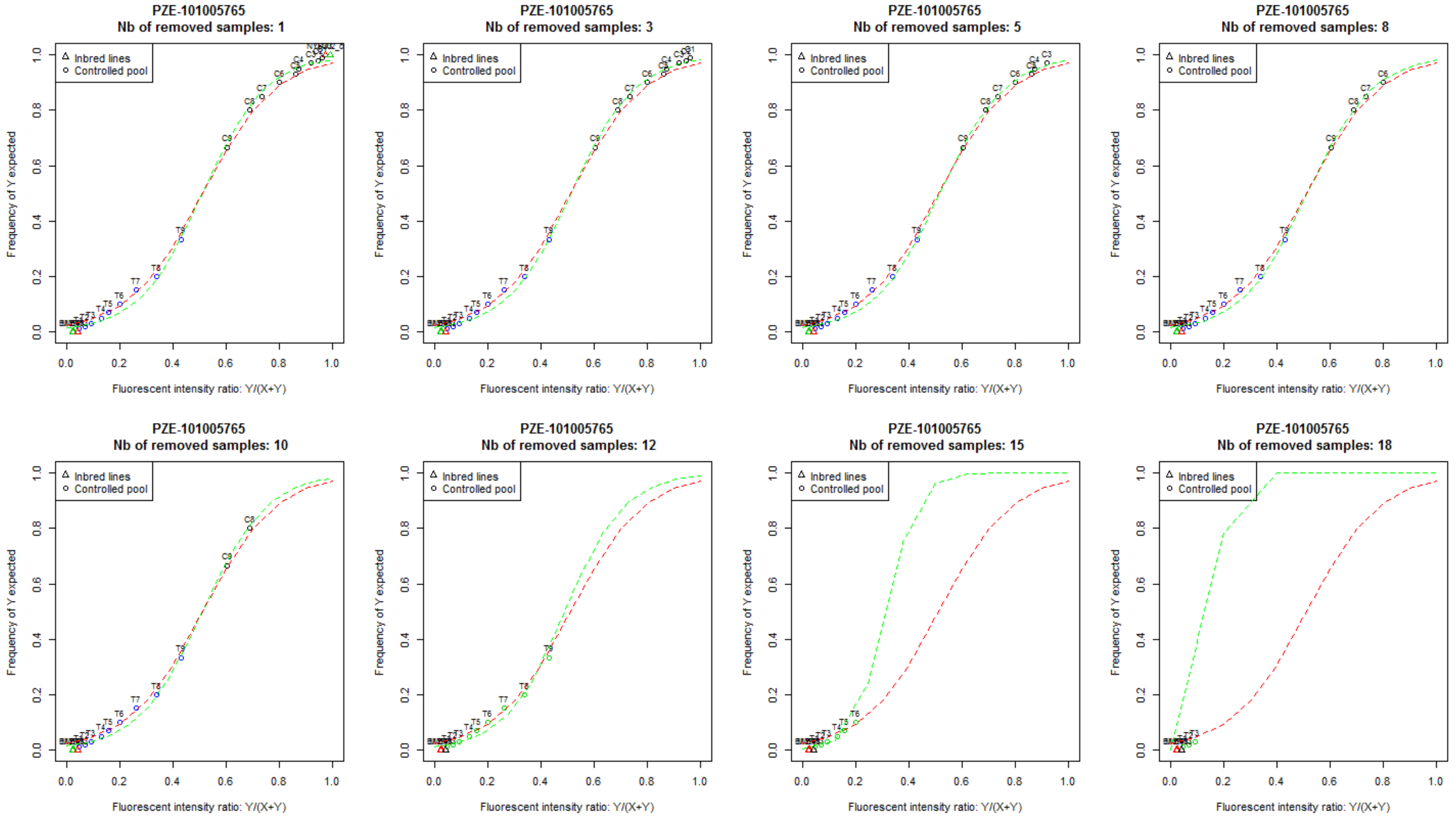


Figure S4: Effect of allelic frequency range on the prediction of allelic frequency in controlled pools for SNP PZE-101005765. Each plot represents the relationship between fluorescent intensity ratio (FIR) and known allelic frequency in controlled pools after removing 1, 3, 5, 8, 10, 12, 15, 18 samples with the highest allelic frequency. Green dotted lines represent the specific predictive equation established with the remaining samples. . Red dotted lines represent the common predictive equation calibrated on 1000 SNPs selected for maximizing range of allelic frequency in controlled pools. Number of removed samples is indicated above each plot.

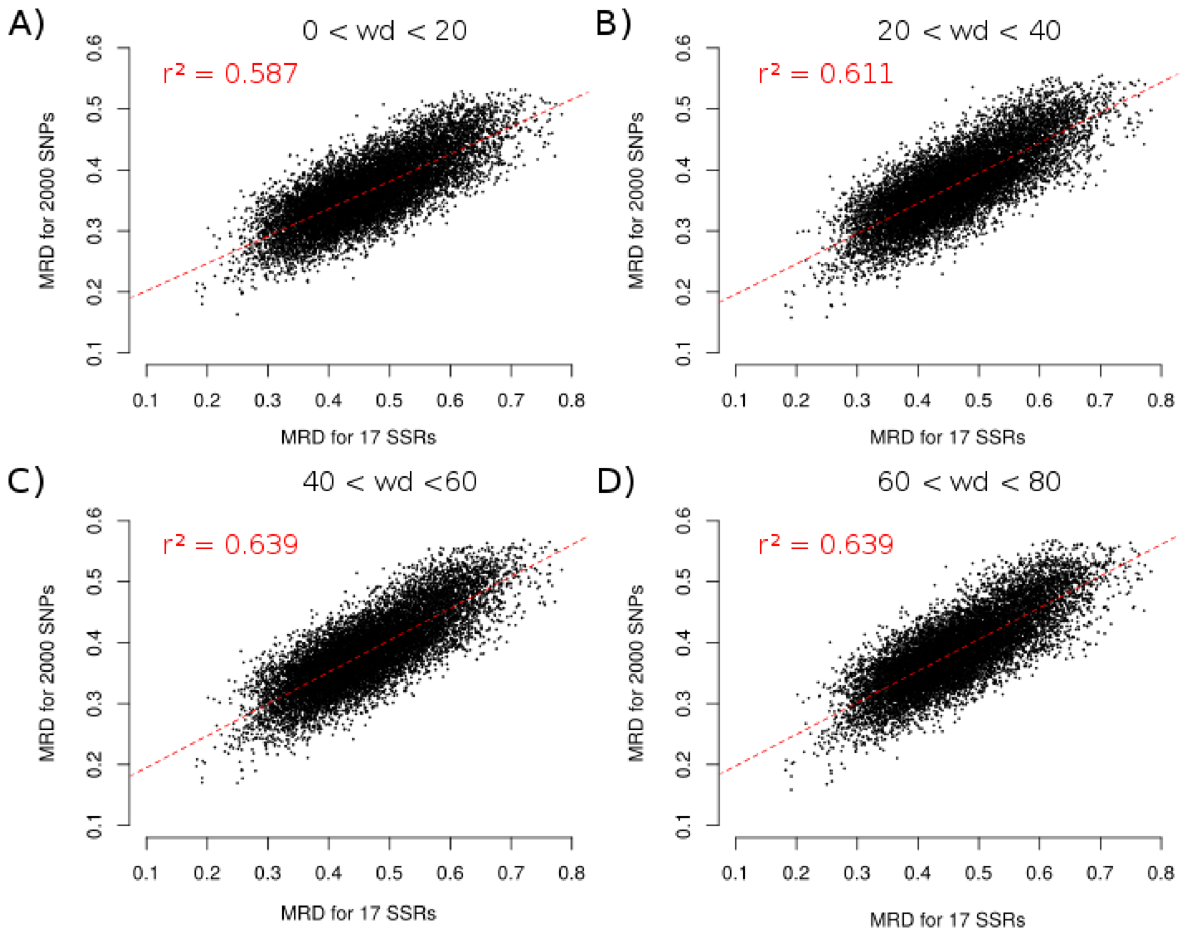


**Figure S5: Correlation between Modified Roger's Distances (MRD) obtained with 17 SSR markers and sets of 2,000 SNPs with different values of weighted deviation criterion (wd). A) 17 SSRs vs. 2,000 SNPs with wd between 0 and 20, B) 17 SSRs vs. 2,000 SNPs with wd between 20 and 40, C) 17 SSRs vs. 2,000 SNPs with wd between 40 and 60, D) 17 SSRs vs. 2000 SNPs with wd between 60 and 80. Red dotted lines represent linear regression between $MRD_{SSR}$ and $MRD_{SNP}$. Coefficient of determination ($r^2$) is reported on top left of each plot. 2000 SNPs were randomly selected within each wd class..**

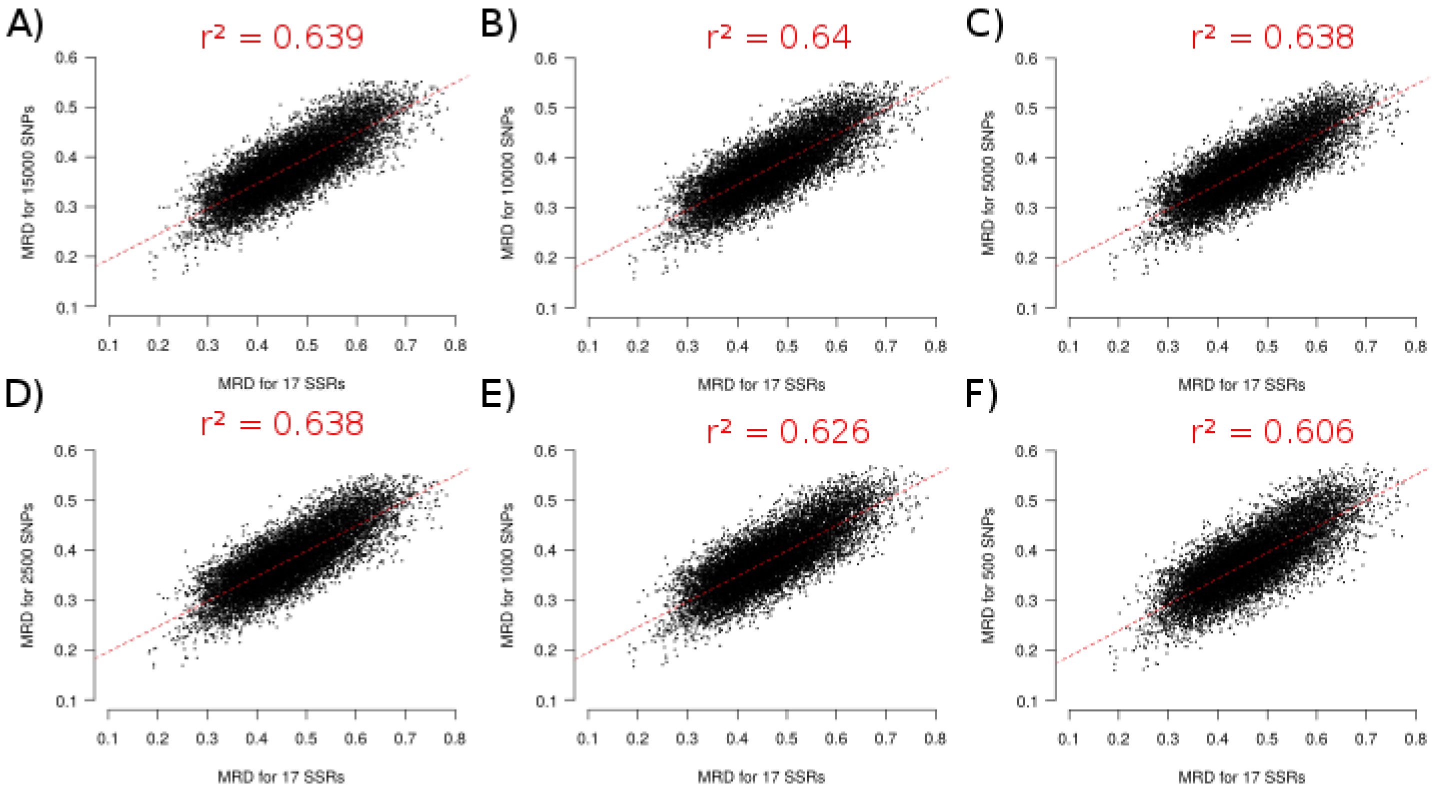


**Figure S6: Effect of SNP number on correlation between Modified Roger's Distances (MRD) obtained with 17 SSR markers and with SNPs. A) 17 SSRs vs 15,000, B) 17 SSRs vs 10,000 SNPs, C) 17 SSRs vs 5,000 SNPs, D) 17 SSRs vs 2500 SNPs E) 17 SSRs vs 1000 SNPs, F) 17 SSRs vs 500. Red dotted lines represents linear regression between $MRD_{SSR}$ and $MRD_{SNP}$. Coefficient of determination ($r^2$) is reported above each plot.**
