## Supplementary Tables 1-3 for "Deciphering the genetic diversity of landraces with high-throughput SNP genotyping of DNA bulks: methodology and application to the maize 50k array"

**Table S1: Mean absolute error (MAE) of allelic frequency prediction estimated by K-fold cross-validation in controlled pools. Logistic regression equations were calibrated on 800 SNPs chosen at random among 1,000 SNPs (training set), then used to predict the remaining 200 SNPs (validation set). Five random K-fold sets of 800/200 SNP were considered. MAE was calculated for each SNP and sample, excluding those with an expected allelic frequency of 0 or 1. MAE was averaged per SNP. Minimum, First Quartile, Median, Mean, Third Quartile, Maximum and standard deviation was estimated across 200 SNPs**

| K-fold set of 200 SNPs | Minimum | First Quartile | Median | Mean | Third Quartile | Maximum | SD* |
|---|---|---|---|---|---|---|---|
| # 1 | 0.0125 | 0.0335 | 0.0507 | 0.0689 | 0.0831 | 0.3954 | 0.0562 |
| # 2 | 0.0161 | 0.0337 | 0.0574 | 0.0764 | 0.0956 | 0.2884 | 0.0591 |
| # 3 | 0.0134 | 0.0313 | 0.0541 | 0.0695 | 0.0919 | 0.2909 | 0.0518 |
| # 4 | 0.0119 | 0.0319 | 0.0524 | 0.0697 | 0.0923 | 0.3019 | 0.0489 |
| # 5 | 0.0136 | 0.0354 | 0.0579 | 0.0747 | 0.0966 | 0.2811 | 0.0529 |
| Average accross K-fold | 0.0135 | 0.0331 | 0.0545 | 0.0719 | 0.0919 | 0.3115 | 0.0538 |

* SD = standard deviation

**Table S2: Mean of absolute error of allele frequency prediction for different expected frequency in two series of controlled pools, including 6 parental inbred lines, for 23,412 SNPs.**

| Expected Allele Frequency | Mean of Absolute Error (MAE) | SD of absolute error | # of observations |
|---|---|---|---|
| 0.000 | 0.000 | 0.005 | 131719 |
| 0.010 | 0.019 | 0.012 | 3495 |
| 0.020 | 0.022 | 0.010 | 3495 |
| 0.030 | 0.021 | 0.016 | 3495 |
| 0.050 | 0.017 | 0.022 | 3495 |
| 0.070 | 0.020 | 0.026 | 3495 |
| 0.100 | 0.029 | 0.036 | 3495 |
| 0.150 | 0.048 | 0.050 | 3495 |
| 0.200 | 0.061 | 0.064 | 3495 |
| 0.333 | 0.090 | 0.080 | 10054 |
| 0.400 | 0.096 | 0.081 | 6559 |
| 0.425 | 0.106 | 0.086 | 6559 |
| 0.450 | 0.108 | 0.086 | 6559 |
| 0.465 | 0.111 | 0.086 | 6559 |
| 0.475 | 0.106 | 0.086 | 6559 |
| 0.485 | 0.113 | 0.088 | 6559 |
| 0.490 | 0.110 | 0.088 | 6559 |
| 0.495 | 0.106 | 0.086 | 6559 |
| 0.505 | 0.107 | 0.085 | 7708 |
| 0.510 | 0.111 | 0.087 | 7708 |
| 0.515 | 0.113 | 0.087 | 7708 |
| 0.525 | 0.106 | 0.085 | 7708 |
| 0.535 | 0.112 | 0.085 | 7708 |
| 0.550 | 0.108 | 0.084 | 7708 |
| 0.575 | 0.108 | 0.084 | 7708 |
| 0.600 | 0.100 | 0.079 | 7708 |
| 0.667 | 0.097 | 0.075 | 11519 |
| 0.800 | 0.064 | 0.056 | 3811 |
| 0.850 | 0.051 | 0.042 | 3811 |
| 0.900 | 0.029 | 0.027 | 3811 |
| 0.930 | 0.018 | 0.020 | 3811 |
| 0.950 | 0.015 | 0.018 | 3811 |
| 0.970 | 0.021 | 0.011 | 3811 |
| 0.980 | 0.023 | 0.007 | 3811 |
| 0.990 | 0.018 | 0.011 | 3811 |
| 1.000 | 0.001 | 0.005 | 178652 |
| NA | NA | NA | 57360 |
| Overall | 0.032 | 0.064 | 561888 |

**Table S3: Allelic frequencies correlation ($r^2$) and pairwise Modified Roger's Distance (MRD) between replicates of a same landrace.**

| Landrace | Sample Replicate 1 | Sample Replicate 2 | $r^2$ | Pairwise MRD |
|---|---|---|---|---|
| Pol3 | WG0109808-DNAH04 | WG0109808-DNAA05 | 0.466 | 0.295 |
| Cze5 | WG0109808-DNAF01 | WG0109808-DNAE01 | 0.947 | 0.098 |
| Hun4 | WG0109808-DNAB04 | WG0109808-DNAC04 | 0.954 | 0.087 |
| Hun2 | WG0109808-DNAH03 | WG0109808-DNAA04 | 0.916 | 0.109 |
| USA10 | WG0109808-DNAG06 | WG0109808-DNAH06 | 0.944 | 0.097 |
| Ita6 | WG0109808-DNAF04 | WG0109808-DNAE04 | 0.948 | 0.092 |
| Sp31 | WG0109808-DNAB02 | WG0109808-DNAC02 | 0.932 | 0.104 |
| Sp36 | WG0109808-DNAD02 | WG0109808-DNAE02 | 0.905 | 0.109 |
| Sp3 | WG0109808-DNAA03 | WG0109808-DNAB03 | 0.897 | 0.120 |
| Sp11 | WG0109808-DNAH02 | WG0109808-DNAG02 | 0.909 | 0.100 |

**Table S4 (.csv): Table of 166 landraces samples corresponding to 156 landraces accessions**

**Table S5 (.csv): Table of 333 inbred lines including 6 parental inbred lines of 2 controlled pools**
